# Preferential IsomiR Enrichment in Extracellular Vesicles Improves Identification of Their Cellular Origins

**DOI:** 10.64898/2026.05.10.724151

**Authors:** Rony Chowdhury Ripan, Xiaoman Li, Haiyan Hu

## Abstract

Extracellular vesicles (EVs) carry microRNAs (miRNAs) that mediate intercellular communication and have strong potential as disease biomarkers, yet the roles of miRNA isoforms (isomiRs) in EVs remain poorly understood. Here, we analyzed 96 human EV and corresponding source samples from nine public datasets. We found that EV samples consistently contained substantially higher proportions of isomiR reads than their corresponding source samples, indicating widespread isomiR enrichment in EVs. Although individual isomiRs showed limited reproducibility across biological replicates and limited sharing between EVs and their corresponding source samples, the parent miRNAs that generated these isomiRs remained highly reproducible across replicates and strongly shared between EV-source pairs. Despite extensive isomiR diversification, EV-source pairs retained highly correlated miRNA expression profiles. Using integrated miRNA- and isomiR-related features, we further developed a random forest model that successfully associated EV samples with their corresponding source samples, with improved performance when isomiR information was included. Together, our results demonstrate that EVs are enriched for biologically meaningful isomiRs while preserving source-associated miRNA landscapes, highlighting the importance of incorporating isomiRs into future EV studies.

## Introduction

Extracellular vesicles (EVs) are membrane-bound particles secreted by virtually all cell types and are now recognized as important mediators of intercellular communication^1,2^. EVs transport diverse molecular cargo, including proteins, lipids, DNA, mRNAs, and small non-coding RNAs, that can influence recipient cell behavior and contribute to processes such as immune regulation, tumor progression, tissue homeostasis, neurodegeneration, and metabolic regulation^3-6^; Because EVs can be isolated from accessible biofluids and retain molecular signatures reflective of their cell origin, they have attracted substantial attention as minimally invasive biomarkers and therapeutic delivery vehicles^5,7,8^. However, EV populations in biofluids are highly heterogeneous and often originate from multiple cell types simultaneously, making it difficult to determine the cellular origin of EV samples without experimental labeling, lineage tracing, or enrichment strategies^9,10^. Consequently, computational approaches capable of associating EVs with their source cells could substantially improve the interpretation of EV datasets and broaden the translational utility of EV-based biomarkers.

Among EV cargo molecules, microRNAs (miRNAs) have attracted substantial attention due to their central role in post-transcriptional gene regulation and their remarkable stability in extracellular environments^11-14^. EV-associated miRNAs can be transferred to recipient cells, where they alter gene expression and modulate diverse biological pathways^3,15^. Numerous studies have implicated EV miRNAs in angiogenesis, immune modulation, metastasis, and metabolic regulation^16,17^. The lipid bilayer of EVs protects miRNAs from extracellular RNases, further increasing their appeal as circulating diagnostic and prognostic biomarkers^12,18^. Consequently, profiling miRNA content in EVs has become a central focus in studies of EV biogenesis, cargo selection, disease progression, and intercellular communication^19,20^. However, selective loading mechanisms can produce substantial differences between EV and source-cell miRNA profiles^21,22^, complicating efforts to infer EV origins solely from miRNA expression patterns. Although correlations between EV and source-cell miRNA profiles have been reported, computational studies directly attempting to identify the cellular origin of EV samples using small RNA signatures remain limited^23^.

An additional and largely overlooked layer of complexity arises from miRNA isoforms, or isomiRs. isomiRs are generated through alternative Drosha and Dicer cleavage, nucleotide additions or trimming, RNA editing, and other post-transcriptional modifications, producing sequence variants that differ from canonical miRNAs at the 5′ end, 3′ end, or internal positions^24-26^. Importantly, sequence variation at the 5′ end can alter the seed region and thereby redirect target specificity, potentially expanding the regulatory repertoire of miRNAs^24,25,27-29^. Increasing evidence suggests that isomiRs are biologically meaningful rather than sequencing artifacts, as they exhibit tissue specificity, developmental regulation, and disease-associated expression patterns^30,31^. Although several studies have reported the presence of isomiRs in EVs and extracellular RNAs^32-34^, most EV studies either collapse isomiRs into canonical miRNAs or do not systematically analyze their abundance, reproducibility, or functional significance. Consequently, the potential contribution of isomiRs to distinguishing EVs from different cellular origins, and to improving computational EV-source association, remains largely unexplored.

In this study, we investigate whether miRNA and isomiR profiles can be used to computationally associate EV samples with their corresponding source samples, with particular emphasis on the contribution of isomiRs to EV-origin identification. We compiled 96 samples from nine human datasets spanning nineteen tissues, cell types, and experimental conditions and performed a comprehensive analysis of miRNA and isomiR abundance, diversity, reproducibility, and expression similarity between EVs and their corresponding source samples. We show that EV samples consistently exhibit a higher proportion of isomiR reads and increased sequence variation, even at the 5′ end, suggesting expanded regulatory potential relative to source samples. Despite these differences, EVs retain significant correlations with their source samples at the miRNA expression level. Building on these observations, we developed a machine learning framework based on a Random Forest classifier using 22 features derived from miRNA and isomiR characteristics to associate EV samples with their origins. Our results demonstrate that EV-source relationships can be inferred with high accuracy and that features involving isomiRs contribute substantially to model performance. Importantly, removing isomiR-related information markedly reduces predictive accuracy, highlighting the critical role of isomiRs in capturing EV identity. To our knowledge, this work represents one of the first systematic computational efforts to associate EVs with their source samples using small RNA profiles and underscores the importance of incorporating isomiR-level information into EV studies.

## Material and Methods

### EV and source samples

To identify datasets containing both EV samples and their corresponding source samples, we searched the Gene Expression Omnibus database using the keyword combination “extracellular vesicle AND miRNA”. We then manually curated the listed human datasets in which miRNA expression had been profiled by next-generation sequencing in both EVs and their matched source samples. In total, 96 samples from nine datasets representing nineteen tissues, cell lines, cell types, or different experimental conditions were collected (Table 1). Among these samples, four EV-source pairs lacked biological replicates (GSE148900 and GSE50429). Our analyses primarily focused on 48 EV-source pairs.

**Table 1.**
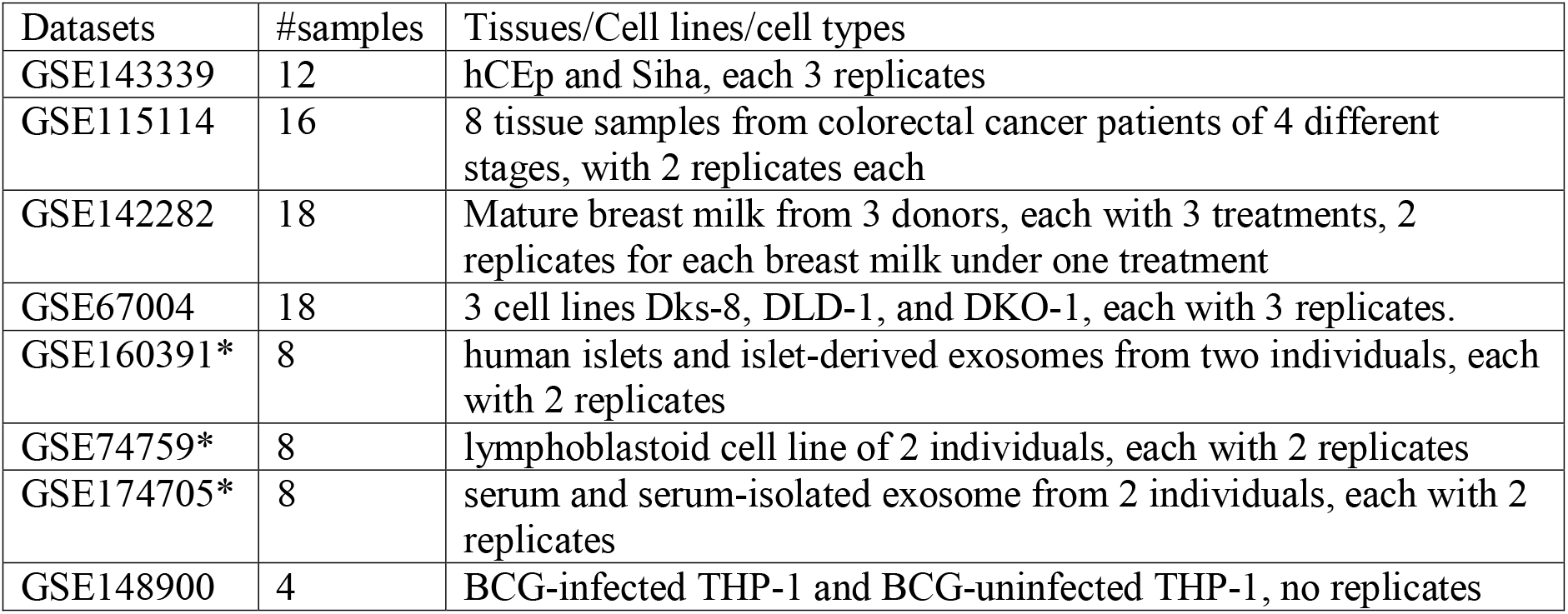
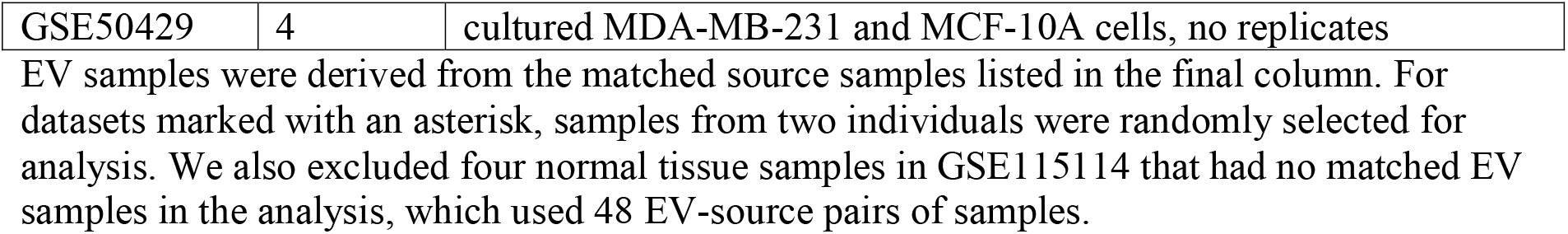
collected EV and source samples.

| Datasets | #samples | Tissues/Cell lines/cell types |
| --- | --- | --- |
| GSE143339 | 12 | hCEp and SiHa, each 3 replicates |
| GSE115114 | 16 | 8 tissue samples from colorectal cancer patients of 4 different stages, with 2 replicates each |
| GSE142282 | 18 | Mature breast milk from 3 donors, each with 3 treatments, 2 replicates for each breast milk under one treatment |
| GSE67004 | 18 | 3 cell lines Dks-8, DLD-1, and DKO-1, each with 3 replicates. |
| GSE160391* | 8 | human islets and islet-derived exosomes from two individuals, each with 2 replicates |
| GSE74759* | 8 | lymphoblastoid cell line of 2 individuals, each with 2 replicates |
| GSE174705* | 8 | serum and serum-isolated exosome from 2 individuals, each with 2 replicates |
| GSE148900 | 4 | BCG-infected THP-1 and BCG-uninfected THP-1, no replicates |
| GSE50429 | 4 | cultured MDA-MB-231 and MCF-10A cells, no replicates |

EV samples were derived from the matched source samples listed in the final column. For datasets marked with an asterisk, samples from two individuals were randomly selected for analysis. We also excluded four normal tissue samples in GSE115114 that had no matched EV samples in the analysis, which used 48 EV-source pairs of samples.

### Data processing

Raw sequencing reads were downloaded from GEO using the Sequence Read Archive Toolkit (fasterq-dump)^35^. Adapter trimming and low-quality read filtering were performed using Trimmomatic^36^. The processed reads were subsequently analyzed using sRNAbench^37^ and miRge3.0^38^. Both tools mapped processed reads to human mature miRNA sequences from miRBase release 22.1^39^ for miRNA and isomiR profiling.

In sRNAbench, miRNA and isomiR detection was performed using the seed-alignment functionality of Bowtie^40^ with a seed length of 20 nucleotides. Reads identical to canonical mature miRNA sequences were classified as canonical miRNAs, whereas reads exhibiting sequence or length variations were classified as isomiRs.

In contrast, miRge3.0 uses a hierarchical alignment strategy. Reads were first aligned to mature miRNAs under stringent exact-match conditions. Unmapped reads were subsequently aligned against hairpin miRNA, noncoding RNA, and coding RNA libraries. Remaining unmapped reads were then realigned to known miRNAs under relaxed conditions, where the first nucleotide and last three nucleotides were ignored and up to three mismatches were allowed to facilitate isomiR detection. To reduce false-positive assignments, miRge3.0 required at least two reads from the stringent alignment step for each detected miRNA species.

For sRNAbench, miRNA and isomiR information was extracted from the microRNAannotation.txt output file. Reads labeled as “canonical” in the isoLabel column were considered miRNA reads, whereas all remaining labels were classified as isomiRs. Mature miRNA annotations were obtained from the matureName column, and expression values were quantified using Reads Per Million (RPM)-normalized values from the RPMTotal column.

For miRge3.0, miRNA and isomiR assignments were obtained from the mapped.csv output file. miRNAs and isomiRs were identified using the “exact miRNA” and “isomiR miRNA” columns, respectively. RPM-normalized miRNA expression values were extracted from the miR.RPM.csv file.

### Training and testing data

We selected a Random Forest classifier to associate EV samples with their source samples because of its robust performance on relatively small datasets, limited hyperparameter tuning requirements, and ability to estimate feature importance^41^.

To construct negative samples, each EV sample was paired with all non-matching source samples, generating 2,256 negative EV-source pairs. From these, 48 negative pairs were randomly selected and combined with the 48 positive pairs to generate a balanced dataset consisting of 96 pairs.

An independent test dataset containing 22 pairs (11 positive and 11 negative) was randomly selected from the above 48 positive and 2256 negative EV-source pairs. Importantly, all replicates associated with a selected EV-source pair were kept entirely within the same dataset partition to prevent data leakage between training and testing. The remaining 74 pairs were used for model training and five-fold cross-validation. To further evaluate model performance under class imbalance, we augmented the independent test dataset with 11 additional negative EV-source pairs randomly selected from the remaining unused negative pairs.

### Features to associate EV with source samples

A total of 22 features were included in the random forest models.

We hypothesized that EV samples retain expression characteristics of their source samples. Therefore, 12 features were designed to quantify similarities in miRNA expression profiles between EV and source samples. The first four features consisted of Pearson’s and Spearman’s correlation coefficients computed using either all detected miRNAs or only miRNAs shared between the EV and source samples. For each correlation metric, two additional contrastive features were calculated: Mean contrastive score, defined as the difference between the correlation of an EV-source pair and the average correlation between the same EV sample and all other source samples; Maximum contrastive score, defined as the difference between the correlation of an EV-source pair and the maximum correlation between this EV sample and any alternative source sample. These contrastive measures quantified whether an EV sample was more similar to its true source than to competing source samples.

We further hypothesized that true EV-source pairs would share more miRNAs and isomiRs than false pairs. Accordingly, six features quantified the overlap of miRNAs and isomiRs between EV and source samples. The first three features were percentage of shared miRNAs, percentage of shared isomiRs, and percentage of shared miRNAs producing isomiRs. Here the percentage was calculated with the denominator as the corresponding miRNAs or isomiRs in both the EV and the source sample. We also had two more features: percentage of EV-unique miRNAs and percentage of EV-unique isomiRs, which had the denominator as the corresponding miRNAs or isomiRs in the EV sample under consideration. The last feature was the percentage of the top 100 highly expressed miRNAs in the source sample that were shared between the EV-source pair, which emphasized the importance of the highly expressed genes instead of any miRNA genes. We did not consider isomiRs here because their expression was difficult to measure without a high sequencing depth.

Four additional features were included to capture complementary aspects of EV-source similarity. The first feature was an enrichment score quantifying the enrichment of source-specific miRNAs within the EV expression profile. Source-specific signatures were generated by calculating log2 fold changes between each source sample and the median expression across all remaining source samples, followed by selecting the top 50 upregulated miRNAs. Enrichment scores were then computed using a running-sum statistic similar to Gene Set Enrichment Analysis^42^. The second feature was the Jaccard similarity between the top 50 source-specific miRNAs and the top 50 highly expressed miRNAs in the EV sample. The third feature was the difference in Gini coefficients between EV and source samples, computed as Gini Difference = Gini_*EV*_ - Gini_*Source*_. This metric quantified differences in miRNA expression inequality between samples. The final feature was cosine similarity computed from normalized expression vectors of shared miRNAs between EV and source samples.

### IsomiR classification

We considered four types of isomiRs: 5’ isomiRs, 3’ isomiRs, both 5’ and 3’ isomiRs, and others. The 5’ isomiRs have variation in the first eight miRNA positions, length or nucleotide variations, while do not have any variation in the last eight miRNA positions. The 3’ isomiRs have variations in the last eight positions, which can be length or nucleotide variations, while do not have any variation in the first eight positions. Both 5’ and 3’ isomiRs have variations in both ends. The fourth type of isomiRs have variations only in the middle of miRNAs.

## Results

### EV samples have much more abundant isomiRs than source samples

We first investigated the abundance of miRNA and isomiR reads in individual EV and source samples. Across the 48 EV samples, both sRNAbench and miRge3.0 consistently detected substantial proportions of reads corresponding to isomiRs (Table 2). Using sRNAbench, isomiR reads accounted for an average of 71.40% of mapped reads in EV samples, whereas miRNA reads accounted for 28.60%. In contrast, miRge3.0 identified an average of 48.13% isomiR reads and 51.87% miRNA reads in EV samples. The lower proportion of isomiR reads identified by miRge3.0 likely reflects its more stringent alignment and filtering strategy for isomiR detection. To reduce potential false-positive isomiR calls arising from sequencing errors or random degradation products, we further restricted analyses to isomiRs supported by at least three identical reads. Under this more stringent criterion, isomiRs still accounted for an average of 43.90% of mapped reads in EV samples, indicating that isomiR enrichment is an important characteristic of EV samples.

**Table 2:**
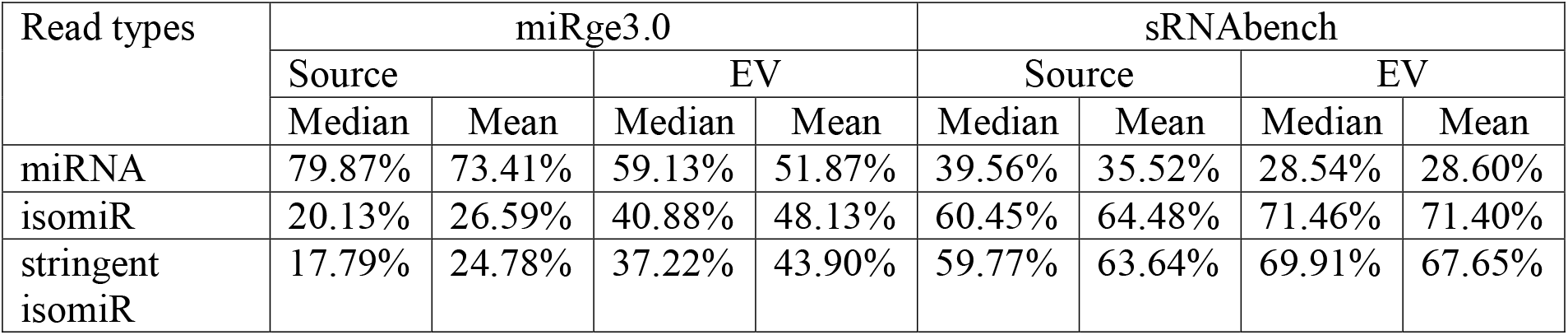
Percentages of reads mapped to miRNA, isomiR and more stringent isomiRs.

| Read types | miRge3.0 |  |  |  | sRNAbench |  |  |  |
| --- | --- | --- | --- | --- | --- | --- | --- | --- |
|  | Source |  | EV |  | Source |  | EV |  |
|  | Median | Mean | Median | Mean | Median | Mean | Median | Mean |
| miRNA | 79.87% | 73.41% | 59.13% | 51.87% | 39.56% | 35.52% | 28.54% | 28.60% |
| isomiR | 20.13% | 26.59% | 40.88% | 48.13% | 60.45% | 64.48% | 71.46% | 71.40% |
| stringent isomiR | 17.79% | 24.78% | 37.22% | 43.90% | 59.77% | 63.64% | 69.91% | 67.65% |

We next examined source samples using the same analytical framework. Similar trends were observed, with sRNAbench detecting more reads assigned to isomiRs than miRNAs, whereas miRge3.0 identified comparable but slightly lower proportions of isomiR reads. Importantly, both tools consistently showed that EV samples contained substantially higher proportions of isomiR reads than their corresponding source samples. For example, sRNAbench detected average isomiR read proportions of 71.40% and 64.48% in EV and source samples, respectively (Table 2). Similarly, miRge3.0 detected average isomiR read proportions of 48.13% in EV samples compared with only 26.59% in source samples (Table 2). This enrichment remained evident under stricter isomiR definitions, suggesting that elevated isomiR abundance may represent an intrinsic feature of EV small RNA content, consistent with previous reports describing abundant extracellular isomiR populations^32,33^.

We further investigated whether highly expressed miRNAs generated larger numbers of detectable isomiRs. Within each sample, miRNAs were ranked according to read abundance, and the number of associated isomiRs was compared between the top and bottom halves of the ranked list using the Mann-Whitney U test. In both EV and source samples, highly expressed miRNAs consistently exhibited significantly larger numbers of isomiRs and isomiR-associated reads than lowly expressed miRNAs (corrected p < 0.01) regardless of whether sRNAbench or miRge3.0 was used. This relationship remained significant when analyses were restricted to miRNAs shared between EV and source samples. In approximately 90% of samples, the same trend was also observed among EV- or source-specific miRNAs, although the remaining samples likely lacked sufficient numbers of unique miRNAs for robust statistical comparison. Overall, these findings suggest that isomiR diversity is strongly associated with miRNA abundance, consistent with previous observations in small RNA sequencing studies^30,31^.

Finally, we classified isomiRs into four categories according to the positions of sequence variation relative to canonical miRNAs: 5′ isomiRs, 3′ isomiRs, both 5′ and 3′ isomiRs, and internal isomiRs (Materials and Methods). In miRge3.0-processed source samples, 7.29% of isomiRs were classified as 5′ isomiRs and 54.45% as both 5′ and 3′ isomiRs, indicating that approximately 61.74% of isomiRs contained 5′-end variations. In EV samples, the proportion of isomiRs containing 5′ variations increased further, with 3.61% classified as 5′ isomiRs and 75.16% classified as both isomiRs, yielding a total of 78.77% of isomiRs with 5′ alterations. sRNAbench produced similar results, detecting 50.46% and 56.25% of isomiRs with 5′ variations in source and EV samples, respectively. Because sequence variation at the 5′ end can alter the miRNA seed region and redirect target specificity^27,28^, the increased prevalence of 5′ isomiRs in EVs may substantially expand their potential regulatory target repertoire.

### Although only a fraction of isomiRs are reproducible between replicates, their corresponding miRNAs remain highly reproducible

To evaluate the reproducibility of miRNAs and isomiRs, we analyzed the 48 replicated EV-source sample pairs. Here, a miRNA or isomiR was considered “shared” if it was detected in both replicate samples. Both sRNAbench and miRge3.0 showed that miRNAs were highly reproducible between replicates, whereas isomiRs exhibited substantially lower reproducibility.

Using miRge3.0, an average of 69.54% of miRNAs (median = 69.76%) were shared between replicated samples. Similarly, sRNAbench detected an average of 70.22% shared miRNAs (median = 71.36%). In contrast, only 26.78% (median = 24.16%) of isomiRs identified by miRge3.0 and 35.20% (median = 34.07%) identified by sRNAbench were shared between replicates. Despite this lower reproducibility at the isomiR sequence level, the corresponding parent miRNAs remained highly consistent. Specifically, more than 66.60% of miRNAs associated with detected isomiRs were shared between replicates according to both tools, with median values exceeding 69.90%. These findings suggest that although individual isomiR sequences may vary considerably between samples, the underlying miRNA populations producing these isomiRs remain stable and reproducible.

We next evaluated the reproducibility of miRNA expression profiles between replicated samples. For these analyses, miRNA expression was defined as the total number of reads mapped to a canonical miRNA together with all reads mapped to its associated isomiRs, thereby generating a compounded miRNA expression measurement. Because individual isomiRs are numerous and often lowly expressed, direct quantification of isomiR expression alone may be unreliable without substantially deeper sequencing coverage. We therefore focused on compounded miRNA expression profiles.

Across replicated EV and source samples, both tools detected strong expression concordance. Pearson’s correlation coefficients averaged greater than 0.70, while Spearman’s correlation coefficients averaged greater than 0.60 (all p < 1 × 10□^21^). These results demonstrate that, despite variability at the individual isomiR sequence level, overall miRNA expression landscapes remain highly reproducible across biological replicates.

### EV-source sample pairs exhibit strongly correlated miRNA expression profiles

Because EV samples contained substantially enriched isomiR populations relative to their source samples, we next investigated the extent to which EVs retained molecular similarities to their corresponding origins. Specifically, we examined the sharing patterns of miRNAs and isomiRs between EV-source pairs and assessed the similarity of their miRNA expression profiles.

We first analyzed the occurrence of shared miRNAs and isomiRs within individual EV-source pairs. Both sRNAbench and miRge3.0 showed that more than 40.00% of miRNAs were shared between EV samples and their corresponding source samples on average. In contrast, the proportion of shared isomiRs was substantially lower, averaging 12.27% according to sRNAbench and 7.40% according to miRge3.0. This reduction likely reflects the substantially greater diversity and sequence variability of isomiRs. Nevertheless, both tools showed that more than 44.60% of miRNAs corresponding to detected isomiRs were shared between EV and source samples. These findings suggest that although EVs contain expanded isomiR diversity, many of these isomiRs originate from the same underlying miRNA populations present in their source samples.

We next examined the reproducibility of shared and unique miRNAs and isomiRs across replicated EV-source pairs. Specifically, we investigated whether miRNAs and isomiRs shared within one EV-source pair were also observed in corresponding replicated EV-source pairs. Both tools consistently demonstrated that more than 74.00% of common miRNAs identified within one EV-source pair were also shared in another replicated pair. Similarly, more than 44.00% of shared isomiRs and approximately 68.00% of their corresponding parent miRNAs were reproducibly detected across replicated EV-source pairs. In addition, more than 18.00% of EV- or source-specific miRNAs and isomiRs identified in one pair were also detected in the corresponding EV or source samples of replicated pairs. Together, these observations indicate that many detected miRNAs and isomiRs represent biologically reproducible signals rather than random sequencing artifacts.

Finally, we assessed the similarity of miRNA expression profiles between EV samples and their corresponding source samples. Using compounded miRNA expression profiles, both sRNAbench and miRge3.0 detected strong correlations between EV-source pairs. Across all pairs, Pearson’s correlation coefficients exceeded 0.74 on average, while Spearman’s correlation coefficients exceeded 0.62 (all p < 1 × 10□^21^). These results indicate that EVs retain substantial components of the miRNA expression programs of their source samples despite selective RNA loading processes reported previously^21^, supporting the feasibility of computationally associating EVs with their origins.

### miRNAs and isomiRs in EV samples enable accurate identification of their source samples

The analyses above demonstrated that EV samples differ from their source samples through substantial enrichment of isomiRs, while simultaneously retaining correlated miRNA expression patterns. We therefore investigated whether these molecular features could be used to computationally identify the source sample of an EV.

To address this question, we trained a random forest classifier using 22 features derived from miRNA expression correlations, shared miRNA/isomiR content, and global expression similarity metrics (Materials and Methods). The model was trained using 48 positive EV-source pairs and 48 randomly selected negative EV-source pairs.

Across five-fold cross-validation, the full model achieved strong classification performance, with an average accuracy of 0.93, precision of 0.93, recall of 0.95, F1 score of 0.94, and Area Under the Receiver Operating Characteristic Curve (AUROC) of 0.96 on validation datasets (Table 3). Performance remained robust on the independent test dataset (Material and Methods), where the model achieved average precision and recall values of 0.90 and 0.82 respectively. To further evaluate model robustness under more realistic class imbalance conditions, additional negative EV-source pairs were introduced into the independent dataset. Under this setting, the model maintained an AUROC of 0.92 while preserving average precision and recall values of 0.90 and 0.82 respectively, indicating that the classifier generalized well beyond the balanced training scenario.

**Table 3.** Average five-fold performance on different test datasets.

|  | Validation data |  | Independent test data |  | Independent test data with more negatives |  |
| --- | --- | --- | --- | --- | --- | --- |
|  | Full model | miRNA only model | Full Model | miRNA only Model | Full Model | miRNA only Model |
| Accuracy | 0.93 | 0.91 | 0.86 | 0.85 | 0.91 | 0.90 |
| F1 | 0.94 | 0.91 | 0.86 | 0.85 | 0.86 | 0.85 |
| Precision | 0.93 | 0.93 | 0.90 | 0.90 | 0.90 | 0.90 |
| Recall | 0.95 | 0.90 | 0.82 | 0.80 | 0.82 | 0.80 |
| AUROC | 0.96 | 0.97 | 0.87 | 0.88 | 0.92 | 0.92 |

We next examined feature importance of identifying the molecular characteristics most strongly contributing to EV-source association. The five highest-ranked feature categories were: (i) contrastive Pearson correlation of compounded miRNA expression profiles, (ii) cosine similarity, (iii) Jaccard similarity, (iv) contrastive Spearman correlation of compounded miRNA expression profiles, and (v) the proportion of highly expressed source miRNAs shared with the EV sample. Importantly, compounded miRNA expression incorporated reads derived from both canonical miRNAs and associated isomiRs, indicating that isomiR information may contribute to several of the most informative features.

**Fig 1.**
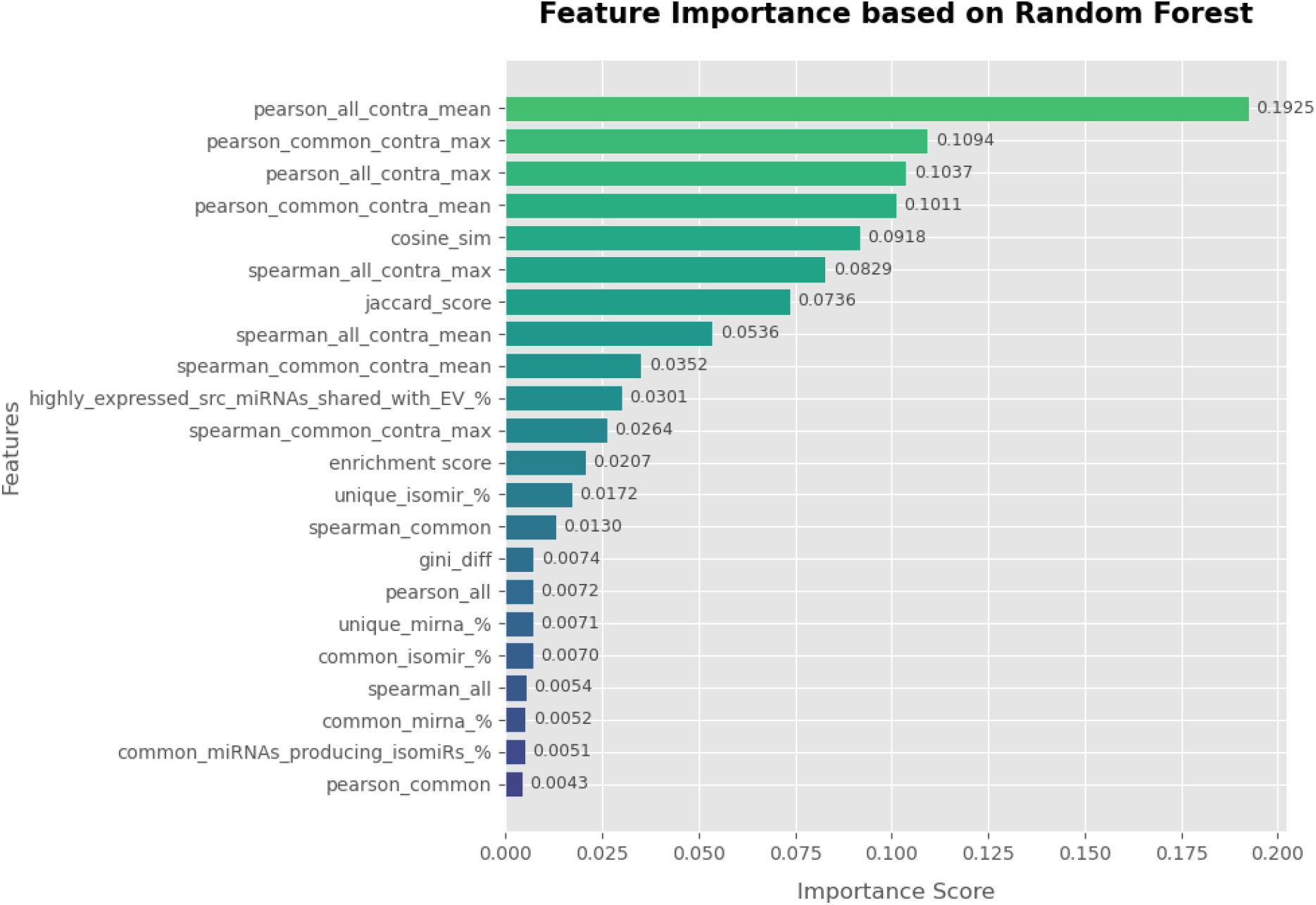
Ranking of features based on Random Forest classifier

To further assess the contribution of isomiRs, we constructed an additional “miRNA-only” model by excluding all isomiR-derived features and removing isomiR-associated reads from miRNA expression quantification. Using identical training and testing procedures, the miRNA-only model consistently demonstrated reduced performance relative to the full model. In the cross-validation dataset, both accuracy and F1 decreased to 0.91 and recall decreased from 0.95 to 0.90. Similar reductions were observed in the independent test dataset and in the imbalanced independent dataset containing additional negative pairs (Table 3). These findings demonstrate that isomiR-associated information improves the ability to computationally associate EV samples with their corresponding source samples.

**Fig 2.**
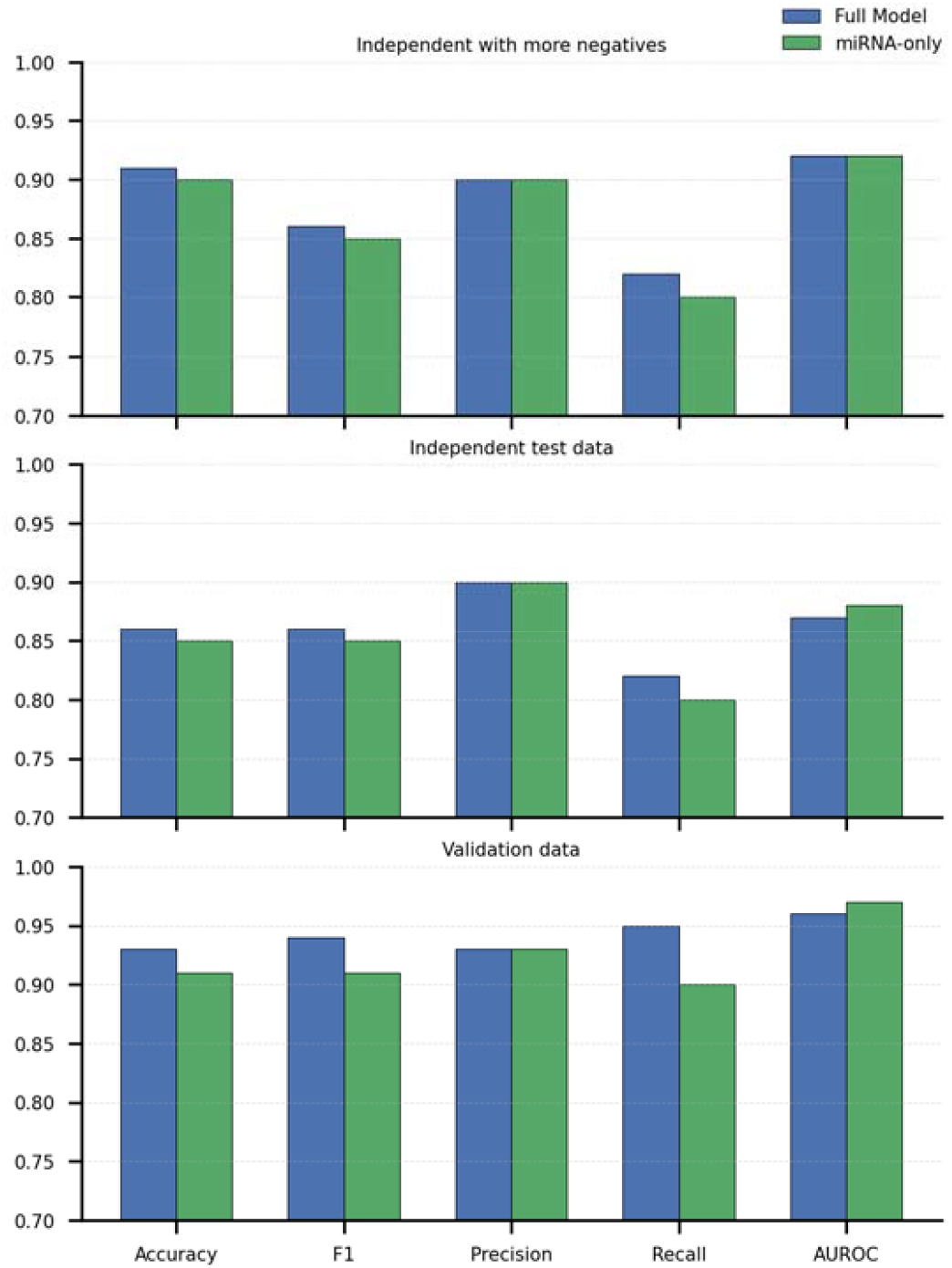
Bar chart comparison of Full model and miRNA only model for different test datasets

Importantly, directly identifying the cellular origin of EVs using computational approaches alone remains a challenging and relatively underexplored problem. Most previous studies have focused on experimentally validating EV origins or characterizing tissue-specific EV biomarkers rather than systematically predicting EV-source relationships using machine learning approaches. Our results therefore suggest that integrating canonical miRNA and isomiR information may provide a promising framework for computationally inferring EV origins from small RNA sequencing data.

## Discussion

In this study, we performed a comprehensive analysis of miRNAs and isomiRs in EV samples and their corresponding source samples across multiple human tissues, cell lines, and biological conditions. By integrating small RNA sequencing datasets from nine independent studies, we demonstrated several key findings. First, EV samples consistently contained substantially higher proportions of isomiR reads than their corresponding source samples. Second, although individual isomiRs showed relatively limited reproducibility across biological replicates, their corresponding parent miRNAs remained highly reproducible, suggesting that isomiR variability occurs within stable miRNA regulatory frameworks. Third, EV samples retained strong miRNA expression correlations with their source samples despite extensive isomiR diversification. Finally, using a machine learning framework integrating miRNA- and isomiR-derived features, we showed that EV samples could be computationally associated with their corresponding source samples with high accuracy. Importantly, models incorporating isomiR information consistently outperformed miRNA-only models, demonstrating that isomiRs contribute biologically meaningful information for EV-source association.

One of the most notable findings of this study is the substantial enrichment of isomiRs within EV samples compared with their source samples. The enrichment of isomiRs in EVs raises the possibility that selective RNA loading mechanisms preferentially package specific miRNA variants into EVs rather than randomly exporting small RNAs from cells. Such selective enrichment may reflect regulated intercellular communication processes, cellular stress responses, or mechanisms for fine-tuning extracellular signaling. Together, these findings support the growing view that isomiRs represent an important and previously underappreciated component of EV biology.

Although both sRNAbench and miRge3.0 consistently supported the major conclusions of this study, the two tools differed substantially in their reported proportions of canonical miRNA and isomiR reads. These differences likely arise from their distinct alignment strategies and criteria for isomiR detection. sRNAbench uses a more permissive alignment framework that allows mismatches near the 3′ end during initial mapping, potentially increasing sensitivity for detecting diverse isomiR species. In contrast, miRge3.0 applies a hierarchical alignment strategy with stringent initial exact matching followed by more conservative relaxed alignment steps, likely reducing false-positive isomiR assignments while sacrificing sensitivity. Despite these methodological differences, both tools consistently identified increased isomiR abundance in EV samples, strong reproducibility of parent miRNA populations, and correlated miRNA expression profiles between EV and source samples. The agreement between two analytically distinct pipelines therefore strengthens the robustness of our conclusions and suggests that the observed biological patterns are unlikely to be artifacts of a particular computational method.

Several limitations of this study should be acknowledged. First, despite integrating datasets from multiple studies, the total number of EV-source pairs remained relatively limited for machine learning applications. Larger and more diverse datasets spanning additional tissues, disease states, and EV isolation methods will likely improve the generalizability and robustness of computational EV-source association models. Second, accurate quantification of individual isomiR expression remains challenging because of the enormous diversity and often low abundance of isomiR species. Deeper sequencing coverage and improved statistical models for low-count small RNAs may enable more reliable isomiR-level expression analyses in future studies. Third, the current study focused primarily on sequence-based and expression-based features derived from miRNAs and isomiRs. Future work could integrate additional molecular layers, including mRNA, long non-coding RNA, protein, lipid, and epitranscriptomic profiles, to further improve EV origin prediction. Moreover, advances in single-EV sequencing technologies and spatially resolved extracellular RNA profiling may provide opportunities to study EV heterogeneity and intercellular communication at substantially higher resolution. Finally, experimental validation of predicted EV-source associations will be important for confirming the biological relevance of computational predictions and for translating these approaches into clinical or diagnostic applications.

## Acknowledgement

This work has been supported by the National Science Foundation [Grant 2514869].

